# Escape of SARS-CoV-2 variants KP1.1, LB.1 and KP3.3 from approved monoclonal antibodies

**DOI:** 10.1101/2024.08.20.608835

**Authors:** Delphine Planas, Isabelle Staropoli, Cyril Planchais, Emilie Yab, Banujaa Jeyarajah, Yannis Rahou, Matthieu Prot, Florence Guivel-Benhassine, Frederic Lemoine, Vincent Enouf, Etienne Simon-Loriere, Hugo Mouquet, Marie-Anne Rameix-Welti, Olivier Schwartz

## Abstract

First-generation anti-SARS-CoV-2 monoclonal antibodies (mAbs) used for prophylaxis or therapeutic purposes in immunocompromised patients have been withdrawn because of the emergence of resistant Omicron variants. In 2024, two novel mAbs, Pemivibart and Sipavibart, have been approved by health authorities, but their activity against contemporary JN.1 sublineages is poorly characterized. We isolated authentic JN.1.1, KP1.1, LB.1 and KP3.3 viruses and evaluated their sensitivity to neutralization by these mAbs in two target cell lines. Compared to ancestral strains, Pemivibart remained moderately active against JN.1 sub-variants, with a strong increase of 50% Inhibitory Concentration (IC50), reaching up to 3 to 15 μg/ml for KP3.3. Sipavibart neutralized JN.1.1 but lost antiviral efficacy against KP1.1, LB.1 and KP3.3. Our results highlight the need for a close clinical monitoring of Pemivibart and raise concerns about the clinical efficacy of Sipavibart.

## INTRODUCTION

The JN.1 lineage arose in late 2023 and rapidly outcompeted previous SARS-CoV-2 variants^1^. Since then, JN.1 continued its evolution, with the appearance of sub-lineages carrying convergent mutations in the Spike (S) protein, notably F456L or R346T, and more recently S31del^2 3^. Sub-lineage specific mutations also appeared, such as Q493E. As of August 2024, the KP.1, KP.2, LB.1 and KP.3 variants, that carry various combinations of these substitutions, represented about 80 % of sequenced circulating strains (Fig. 1A and Fig. S1-S2). These mutations are collectively responsible for increased immune escape from previously infected and vaccinated populations^2 3^.

**Figure 1.**
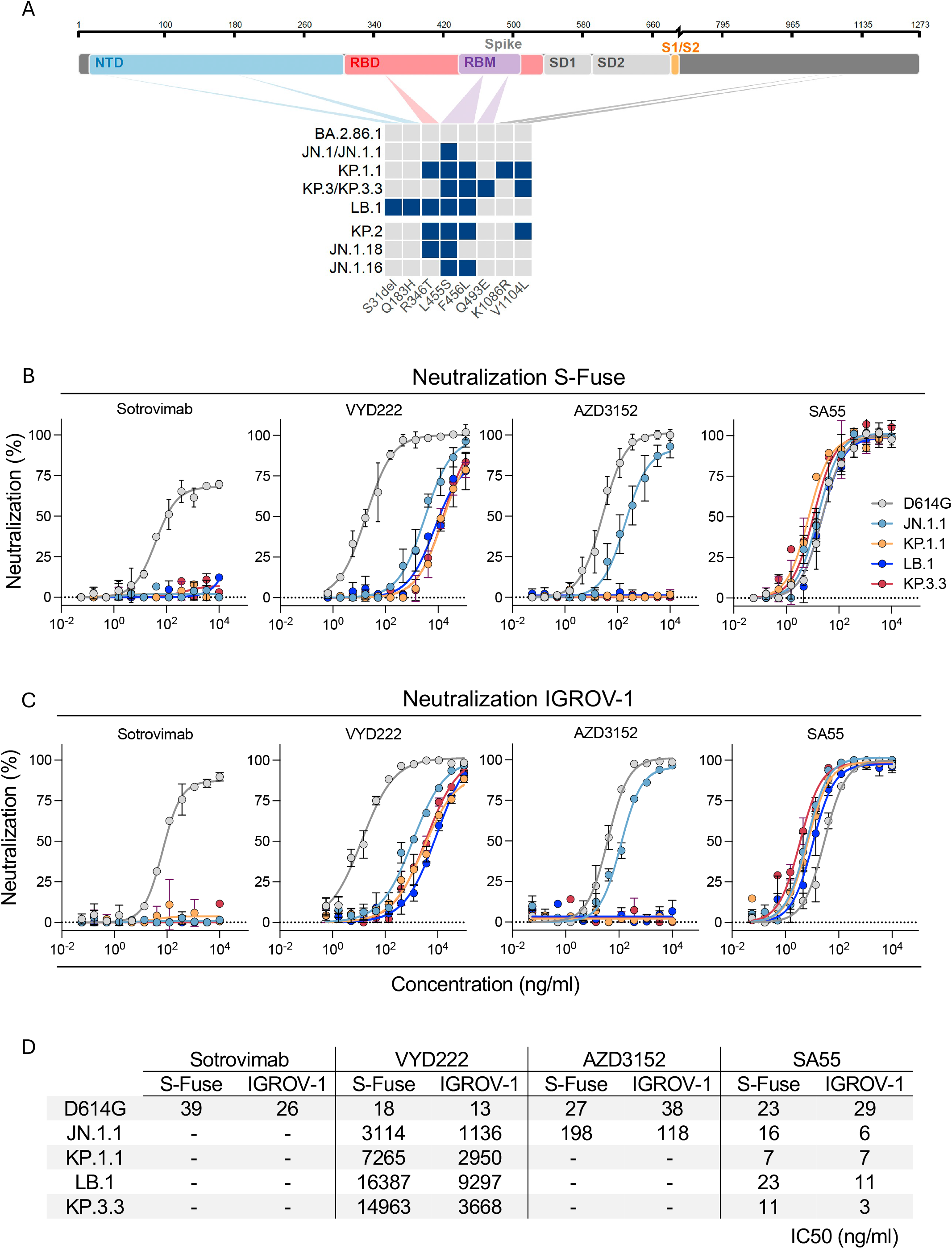
legend. Sequence of SARS-CoV-2 variants and neutralizing activity of mAbs. A. Spike mutations of SARS-CoV-2 variants relative to the spike domains of the BA.2.86.1 parental strain. JN.1, KP.1.1, LB.1, and KP.3.3 were further studied. The mutations in KP.2, JN.1.18 and JN.1.16 are also indicated. B. Neutralization curves of mAbs in S-Fuse cells. Dose-response analysis of neutralization of the indicated variants by Sotrovimab, VYD222, AZD3152, and SA55. Data are presented as mean ± standard deviation of 2-3 independent experiments. C. Neutralization curves of mAbs in IGROV-1 cells. Dose-response analysis of neutralization of the indicated variants by Sotrovimab, VYD222, AZD3152, and SA55. Data are presented as mean ± standard deviation of 2-3 independent experiments. D. EC50 values (in ng/ml) for each mAb against the indicated viral strains in the two cell lines. “-“: no antiviral activity.

The sensitivity of KP.1, LB.1 and KP.3 to monoclonal antibodies (mAbs) developed for clinical use is poorly characterized. First-generation anti-S mAbs, previously approved by the Food and Drug Administration (FDA), the European Medicines Agency (EMA) or other agencies, had their authorization withdrawn after the emergence of Omicron variants, because of escape mutations in the receptor binding domain (RBD) of S. In 2024, novel mAbs, efficient against Omicron variants, have been tested in clinical trials or are available for clinical use in some countries^4^. These include AZD3152, VYD222, and SA55 that belong to different anti-RBD antibody classes and target distinct epitopes. AZD3152 (or Sipavibart™)^5^ was authorized in July 2024 by the EMA, for pre-exposure prophylaxis in patients with immunocompromising conditions and at high risk of developing severe COVID-19^6^. However, AZD3152-resistant viruses, carrying escape mutations at RBD positions 415, 456 and 458, have been described in pre-JN.1 lineages^5^. VYD222/Pemivibart (or Pemgarda™) obtained an emergency use authorization by the FDA in March 2024, for pre-exposure prophylaxis in patients who are moderately to severely immunocompromised^7^. VYD222 preserved *in vitro* efficacy against pre-JN.1 strains carrying the F456L mutation^7^. SA55 was isolated from a SARS-CoV-1 infected, SARS-CoV-2 vaccinated individual and displays a broad sarbecovirus neutralization profile, including JN.1, when used alone or in combination with another mAb (SA58) ^8 9^. SA55 has been tested in a clinical trial initiated in 2023 in China, in patients with hematological disorders who are persistently positive for SARS-CoV-2 ^10^.

Here, we isolated the main SARS-CoV-2 variants circulating in mid-2024 and tested their sensitivity to neutralization by a panel of mAbs.

## METHODS

### Virus isolation

Viral strains were amplified through one or two passages on Vero E6 TMPRSS2 cells and one passage on IGROV-1 cells. Cells were plated in T75 flasks and cultivated in culture media (Dulbecco’s Modified Eagle Medium (DMEM), 10% fetal calf serum, and 1% Penicillin/Streptomycin) at 37°C, 5% CO2. Supernatants were harvested two or three days after viral exposure. Viral supernatants were sequenced directly from nasopharyngeal swabs and after isolation and amplification on IGROV-1 cells to confirm identity, the presence of specific mutations in the spike protein, and the absence of cell culture-derived mutations. The titration of viral stocks was performed on S-Fuse cells ^11 12 1^.

The D614G and JN.1 strains have been described ^13 1^. The KP.1.1 (hCoV-19/France/IDF-RELAB-IPP05044/2024), LB.1 (hCoV-19/France/GES-RELAB-IPP04736/2024), and KP.3.3 strains (hCoV-19/France/BFC-IPP06087/2024) were isolated and amplified by the National Reference Center for Respiratory Viruses hosted by Institut Pasteur.

### Monoclonal antibodies

Sotrovimab (S309) was previously described ^14^. Codon-optimized synthetic DNA fragments coding for the immunoglobulin variable domains of SA55 (BD55-5514) ^8^, AZ3152/Sipavibart ^5^ and VYD222/Pemivibart were synthetized (GeneArt, Thermo Fisher Scientific), and cloned into human IgG1 expression vectors as previously described ^15^. Recombinant IgG1 antibodies were produced by transient co-transfection of Freestyle^™^ 293-F suspension cells (Thermo Fisher Scientific) using PEI-precipitation method and purified from culture supernatants by affinity chromatography using Protein G Sepharose® 4 Fast Flow (GE Healthcare) as previously described ^15^.

### Cell lines

IGROV-1 and S-Fuse (U20S) cells were previously described ^1 11^. Cells were regularly tested negative for mycoplasma.

### Virus titration for neutralizing assay

Titration of viral stocks was performed on S-Fuse and IGROV-1 cells. Neutralization assays were conducted using a multiplicity of infection sufficient to produce about 200 syncytia/well with S-Fuse cells and achieve 40% of infected IGROV-1 cells.

### S-Fuse neutralization assay

U2OS-ACE2 GFP1-10 and GFP11 cells, also termed S-Fuse cells, become GFP+ when they are productively infected by SARS-CoV-2 ^11,13^. Cells were mixed (ratio 1:1) and plated overnight at 12 × 10^3^ per well in a μClear 96-well plate (Greiner Bio-One). The indicated SARS-CoV-2 strains were incubated with serially diluted monoclonal antibodies (mAbs) for 15 min at room temperature and added to S-Fuse cells. Eighteen hours later, cells were fixed with 2% PFA (Electron Microscopy Sciences, cat# 15714-S), washed, and stained with Hoechst (dilution of 1:1,000, Invitrogen, cat# H3570). Images were acquired using an Opera Phenix high-content confocal microscope (PerkinElmer). The number of GFP syncytia and the number of nuclei were quantified using Harmony software (PerkinElmer). The percentage of neutralization was calculated using the number of syncytia with the following formula: 100 × (1 – (value with mAb – value in ‘non-infected’)/(value in ‘no mAb’ – value in ‘non-infected’)). For each mAb, the half maximal inhibitory concentration (IC50) in ng/ml was calculated with a reconstructed curve using the percentage of neutralization at each concentration.

### IGROV-1 neutralization assay

Sixteen hours before infection, 30 × 10^3^ cells per well were seeded in a μClear black 96-well plate (Greiner Bio-One). The indicated SARS-CoV-2 strains were incubated with serially diluted monoclonal antibodies (mAbs) for 15 min at room temperature and added to IGROV-1 cells. Twenty-four hours later, cells were fixed with 2% PFA (Electron Microscopy Sciences, cat# 15714-S). The cells were then intracellularly stained with anti-SARS-CoV-2 nucleoprotein (N) antibody NCP-1 (0.1 μg/mL) as described^1^. The staining was carried out in PBS with 0.05% saponin 1% BSA, and 0.05% sodium azide for 1 h. Cells were then washed twice with PBS and stained with anti-IgG Alexa Fluor 488 (dilution 1:500, Invitrogen; cat# A11029) for 30 minutes before being washed twice with PBS. Hoechst 33342 (Invitrogen, cat# H3570) was added during the final PBS wash. Images were captured using an Opera Phenix high-content confocal microscope (PerkinElmer). The N-positive area and the number of nuclei were quantified using Harmony Software v4.9 (PerkinElmer). The percentage of neutralization was calculated using the N-positive area with the following formula: 100 × (1 – (value with mAb – value in ‘non-infected’)/(value in ‘no mAb’ – value in ‘non-infected’)). For each mAb, the half maximal inhibitory concentration (IC50) in ng/ml was calculated with a reconstructed curve using the percentage of neutralization at each concentration.

### Statistical analysis

Figures were generated using Prism 9 (GraphPad Software). Statistical analysis was conducted using GraphPad Prism 9. Data are mean ±SD of three independent experiments.

### Lineage monitoring

To visualize the evolution of the frequency of SARS-CoV-2 lineages, we analyzed the viral genomic surveillance data deposited in the GISAID database (https://www.gisaid.org; metadata downloaded on July 7^th^, 2024) ^16 17^. The hierarchical relationships between lineages were retrieved from the pangolin GitHub repository (https://github.com/cov-lineages/pango-designation). We analyzed SARS-CoV-2 data collected from January 1^st^, 2024, toAugust 4, 2024, using R 4.3 and ggplot 3.4.3. Mutations that are common and specific to lineages of interest were computed using the outbreak.info R package (https://outbreak-info.github.io/R-outbreak-info) ^18^.

## Data availability

All data supporting the findings of this study are available within the article or from the corresponding author upon reasonable request without any restrictions. The sequencing data generated in this study have been deposited in the GISAID EpiCoV database.

## RESULTS AND DISCUSSION

We examined the sensitivity of SARS-CoV-2 variants JN.1.1, KP.1.1, LB.1 and KP.3.3 to VYD222/Pemivibart, AZD3152/Sipavibart and SA55. We included the ancestral D614G strain as control. We isolated KP.1.1 (which carries the same S as KP.1), LB.1 and KP.3.3 (which carries the same S as KP.3) variants from nasal swabs of individuals with sequence-diagnosed infections. Sequences of outgrown viruses confirmed the identity of the variants (Fig. 1A and Fig. S1-S2). The mAbs were not commercially available for research purposes. Therefore, we retrieved their sequences from public databases and produced biosimilar molecules. As additional control, we used Sotrovimab™ that neutralizes several Omicron strains but not JN.1, and is no longer approved^1^.

We measured the sensitivity of the viral isolates to mAbs using first S-Fuse cells as targets^12^. These cells were engineered to express ACE2 and are thus sensitive to SARS-CoV-2^1, 12^. The four mAbs efficiently neutralized D614G (Fig. 1B), with EC50s of 18-39 ng/ml, corresponding to those described in the literature. As expected, Sotrovimab lost any activity against the four JN.1-derived strains. AZD3152/Sipavibart inhibited JN.1.1, with an EC50 of 198 ng/ml, but no longer neutralized KP.1.1, LB.1 and KP.3.3 (Fig. 1B). The F456L substitution present in the three variants likely mediates this resistance. VYD222/Pemivibart was poorly active against JN.1.1 and displayed a decreased antiviral activity against KP.1.1, LB.1 and KP.3.3 (Fig. 1B). The EC50s reached up to 16,000 ng/ml, corresponding to up to 888-fold reduction of potency against the four variants compared to D614G. The antiviral activity of SA55 was preserved against the variants, with EC50s that remained remarkably low (7 to 23 ng/ml) (Fig. 1B).

We did not isolate a KP.2 variant, but its profile of resistance is likely similar to KP.1, since their S only differ at position 1086, outside of the RBD (Fig. 1A).

We then sought to confirm these results using another cell line. We selected IGROV-1 cells, because they naturally express ACE2 and are highly sensitive to SARS-CoV-2, including Omicron and JN.1 variants^1^. The profile of neutralization of the five SARS-CoV-2 strains was similar in S-Fuse (Fig. 1B) and IGROV-1 cells (Fig. 1C). The IC50 were also in the same range in the two cell types (Fig. 1D).

Altogether, our results indicate that AZD3152/Sipavibart totally lost antiviral activity against the prevalent strains circulating in mid-2024, most likely because of the presence of the F456L substitution in S. Pemivibart remains active against JN.1.1, KP.1.1, LB.1 and KP.3.3, with however a strong increase in IC50. The loss of activity of Pemivibart has been recently reported in a preprint, using VSV-based pseudotypes ^19^. As of August 2024, the KP3.1.1 variant, that combines the F456L and Q493E mutations found in KP.3 and KP.3.3, with the S31 deletion found in LB.1, has been on the rise ^19^. Future work will help assessing the sensitivity of the rapidly diversifying JN.1 family to these mAbs.

Our *in vitro* results may not directly translate into clinical efficacy, but raise concerns about the medical use of Sipavibart, and warrant a close surveillance of Pemivibart, when most of the circulating strains totally or partially escape neutralization by the two antibodies. The mAb SA55 represents a promising alternative.

## ACKNOWLEDGEMENTS

The authors thank Timothée Bruel for critical reading of the manuscript, members of the Virus and Immunity Unit and other teams for discussion and help, Nathalie Aulner and the staff at the UtechS Photonic BioImaging (UPBI) core facility (Institut Pasteur), a member of the France BioImaging network, for image acquisition and analysis, the members of the National Reference Center for Viruses of Respiratory Infections hosted at Institut Pasteur, the RELAB network and the Mutualized Platform for Microbiology (P2M). This work has used the computational and storage services (Maestro cluster) provided by the IT department at Institut Pasteur, Paris.

Work in OS lab is funded by Institut Pasteur, Urgence COVID-19 Fundraising Campaign of Institut Pasteur, Fondation pour la Recherche Médicale (FRM), ANRS, the Vaccine Research Institute (VRI) (ANR-10-LABX-77), Labex IBEID (ANR-10-LABX-62-IBEID), the HERA Project DURABLE (grant 101102733) and LEAPS, DP is supported by the VRI.

## AUTHOR CONTRIBUTIONS

Experimental strategy design, experiments: DP, IS, CP, FG-B, ES-L, HM, M-ARW, OS.

Vital materials: CP, EY, BJ, YR, VE, HM, M-ARW.

Phylogenetic analysis: ES-L

Viral sequencing: EY, BJ, YR, VE, MP, FL, ES-L, M-ARW.

Manuscript writing and editing: DP, ES-L, HM, M-ARW, OS.

