## Supplementary material for "Escape of SARS-CoV-2 variants KP1.1, LB.1 and KP3.3 from approved monoclonal antibodies": Suppl Figures

### Supplementary Figures

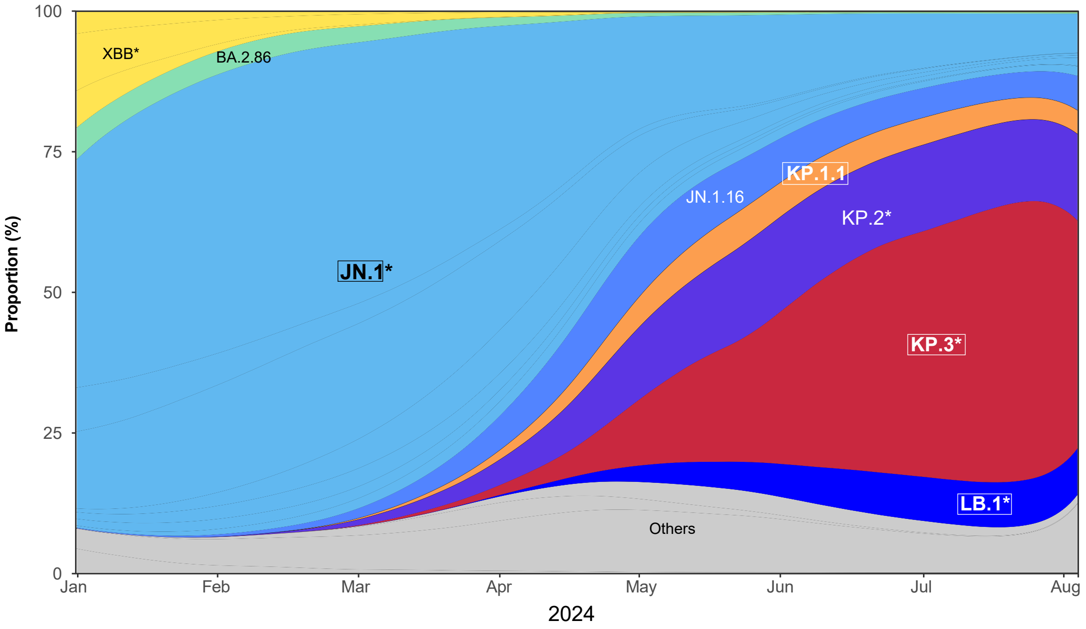


**Figure S1.**

Evolution of the prevalence of the main SARS-CoV-2 lineages from January to August 4, 2024, showing the expansion of JN.1 derived lineages. The star indicates the presence of different sublineages. The variants highlighted with a frame are analyzed in this study.

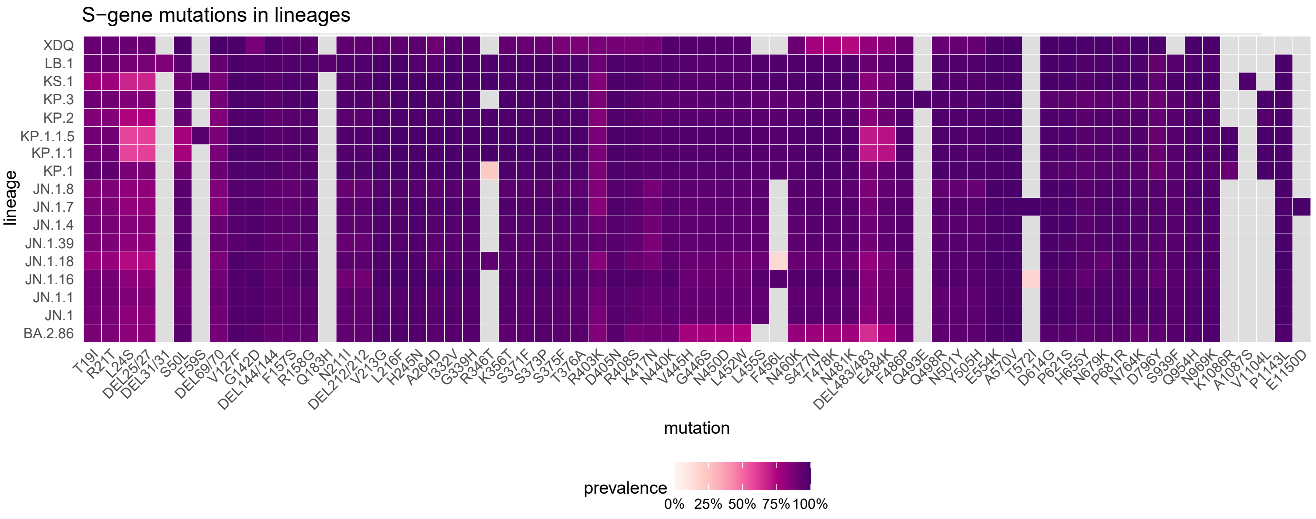


**Figure S2.**

Comparison of spike mutations of selected SARS-CoV-2 variants using Wuhan_Hu-1 (NC_045512) as reference. The color scale reflects the frequency of the mutations within lineages based on the data available on the GISAID epiCOV database.
